# The Composition of Diesel Fuel Influences the Structure of Microbiological Assemblages in Contaminated Storage Tanks

**DOI:** 10.1101/2022.02.09.479836

**Authors:** James G. Floyd, Blake W. Stamps, Caitlin L. Bojanowski, Wendy J. Goodson, Bradley S. Stevenson

## Abstract

Microbiological contamination in diesel storage tanks is widespread and under-reported. The contaminating microorganisms can degrade components of the fuel, which contributes to fouling and corrosion. To better understand the connection between the microorganisms that are responsible for this contamination, the composition of affected fuels, and the resulting impact on fuel quality, we conducted a survey of 106 fuel tanks at 17 military bases across the continental United States. Fuel was sampled from each tank and the resident microbial communities were characterized using high throughput sequencing of small subunit ribosomal RNA gene libraries. Fatty acid methyl esters (FAME) and n-alkanes were characterized and quantified using GC-MS to determine their correlation with the presence of microbial taxa. Redundancy Analyses identified which microbial taxa were more prominent in contaminated fuels. Members of the fungal family *Trichomaceae* were found to be prominent in fuels containing more FAME. Members of the yeast family *Debaryomycetaceae* were found to be prominent in fuels containing more pentadecanoic and oleic acid methyl esters. These relationships between fungal taxa and fuel components were directly tested in growth experiments with representative isolates of the *Trichocomaceae* (Paecilomyces AF001) and *Debaryomycetaceae* (*Wickerhamomyces* SE3) families. *Paecilomyces* was capable of growth on linoleic acid methyl ester but unable to grow on pentadecanoic acid methyl ester, while *Wickerhamomyces* was able to grow on both substrates. Fuel composition may provide some insight into which microorganisms can proliferate but other factors like competition and symbiosis may also drive microbial proliferation, fouling, degradation, and corrosion in diesel fuels.

**Importance:** Biodiesel, widely used as an additive or extender of ultra-low sulfur diesel, can increase the potential for microorganisms to proliferate in storage tanks. It is important to know how the composition of diesel fuels can influence the growth of organisms linked to fuel degradation and microbiologically influenced corrosion. This research describes how certain populations of fungi and bacteria can prevail in fuels of different composition, which can be helpful in predicting biodegradation and biocorrosion, and formulating fuels less susceptible to the growth of problematic organisms.

## Introduction

The U.S. Energy Information Administration predicts that petroleum and other fuel consumption can be expected to rise from the current consumption of 16 million barrels per day to just under 25 million barrels per day by the year 2050 (1). One common transportation fuel is diesel blended with biodiesel (*i*.*e*., biodiesel blends). In recent years, the U.S. Department of Energy has incentivized the use of biodiesel through December 2022 in efforts to offset carbon emissions (2). One common biodiesel blend in the U.S. is B20, containing 20% biodiesel and 80% petroleum diesel. B20 represents a balance of costs, emissions control, cold-weather performance, and engine material compatibility (3). Additionally, biodiesel is readily added to ultralow sulfur diesel (ULSD) at up to 5% v/v to compensate for the loss of lubricity from the desulfurization process (4). Biodiesel (and its’ blends) is also more easily degraded by endogenous environmental microorganisms and has a much higher flash point when compared to ULSD, making it much safer for the environment and those handling the fuel (4–6); However, the high biodegradability of B20 is problematic for operators that store biodiesel containing fuels long term (7).

Biodiesel is composed of fatty acid methyl esters (FAMEs) that are produced through a transesterification reaction with feedstocks of animal, plant, or microbial lipids (8). The FAMEs in biodiesel make the fuel more susceptible to microbial contamination since FAME compounds exist naturally in the environment and are readily degraded by many different microorganisms (9, 10). Additionally, hydrocarbons (*e*.*g*. n-alkanes) in diesel fuel can also be used as an oxidizable substrate for many microorganisms (11). The microbial metabolism of both FAMEs and n-alkanes in fuels manifests itself as microbial contamination within fuel storage infrastructure. Microbial contamination and proliferation in storage tanks containing biodiesel and biodiesel blends leads to increased operating costs through mechanical cleaning of affected storage tanks. Additionally, the contaminating microbial populations can increase corrosion of storage tank materials because their metabolism of the fuel produces acidic metabolic byproducts and their production of biofilms on surfaces cab generate oxygen corrosion cells under aerobic conditions (12). Fungi and bacteria grown on B20 biodiesel can not only degrade FAME and hydrocarbons, but also increase corrosion rates and pitting corrosion of carbon steel (12–14).

All biodiesels or biodiesel blends are not identical. When different feedstocks (*e*.*g*. animal fats, soy, etc.) are used to produce biodiesel, they result in unique FAME profiles with variations in fatty acid chain lengths and degrees of saturation (15). The varying profiles of each biodiesel, therefore, may selection for differing communities of microorganisms that are adapted to preferentially consume specific FAMEs. Microorganisms oxidize FAME using lipases to first hydrolyze the ester bond, converting the FAME to methanol and the corresponding fatty acid, which is then further oxidized to yield ATP (9,16). Different microorganisms have differing capabilities of fatty acid metabolism that depend on fatty acid chain length. Some metabolic pathways even require the use of syntrophic communities to utilize long chain fatty acids (17).

Microorganisms exhibit substrate specificity towards certain FAMEs, which is why each fuel may influence the composition of the microbial community composition within each fuel storage tank.

The major components of most biodiesel blends, except for B100, come from the petroleum itself, which varies in composition based on its provenance and distillation processes used to formulate the final product (18). Diesel fuel primarily consists of hydrocarbons between 7 and 24 carbon atoms in length and is typically comprised of 64% alkanes (including n-, iso-, and cyclo-alkanes), 1-2% alkenes and 34% aromatic hydrocarbons (19, 20). In general, microorganisms preferentially oxidize shorter chained hydrocarbons (21), although there are some exceptions of microorganisms preferentially metabolizing longer chain alkanes (C20+) (22). The combination of compositional differences in diesel fuels and differential metabolic capabilities among microorganisms would represent a highly selective framework that could explain the structure of microbial communities in contaminated fuel.

Microbial community assemblies can shift based on the carbon substrates that are available in the environment (23–26). To determine the relationship between fuel composition and microbial communities in contaminated fuels, we conducted a survey of diesel fuels (ULSD and biodiesel blends) from military installations across the continental United States. Microbial community composition was analyzed via SSU rRNA gene sequencing and fuel composition was analyzed using gas chromatography coupled to mass spectrometry (GC-MS). Previous work identified both fungi and bacteria linked to elevated rates of corrosion when grown on biodiesel and diesel fuels (12, 13, 27). Correlations between fuel substrates and taxa from RDA analyses were tested using the fungal isolates *Paecilomyces* AF001 (Family - *Trichocomaceae*) and *Wickerhamomyces* SE3 (Family -*Debaryomycetaceae*), which have both been previously linked to corrosion risks when grown using biodiesel as a sole carbon and energy source (13). Herein we identify correlative links between the bacterial and fungal microbial communities present within fuel storage tanks across the U.S. and the concentrations of specific FAME and hydrocarbon components of stored fuel. With a better understanding of the relationships between fuel composition and their contaminating microbial communities, we can alert fuel storage operators to the potential risks associated with fouling and microbiologically influenced corrosion for diesel fuel based on its composition.

## Results

### Characterization of n-alkanes and FAME components in fuels

Biodiesel blend and diesel fuels were found to have different profiles of n-alkanes and FAME components. When visualized by PCA, fuels separated by geographic region instead of fuel type (FIG 1A). The largest drivers of alkane variability in the collected fuels were from C9 (nonane)-C20 (eicosane) alkanes and were responsible for much of the separation of samples in the PCA (FIG 1A).

**FIG 1A.**
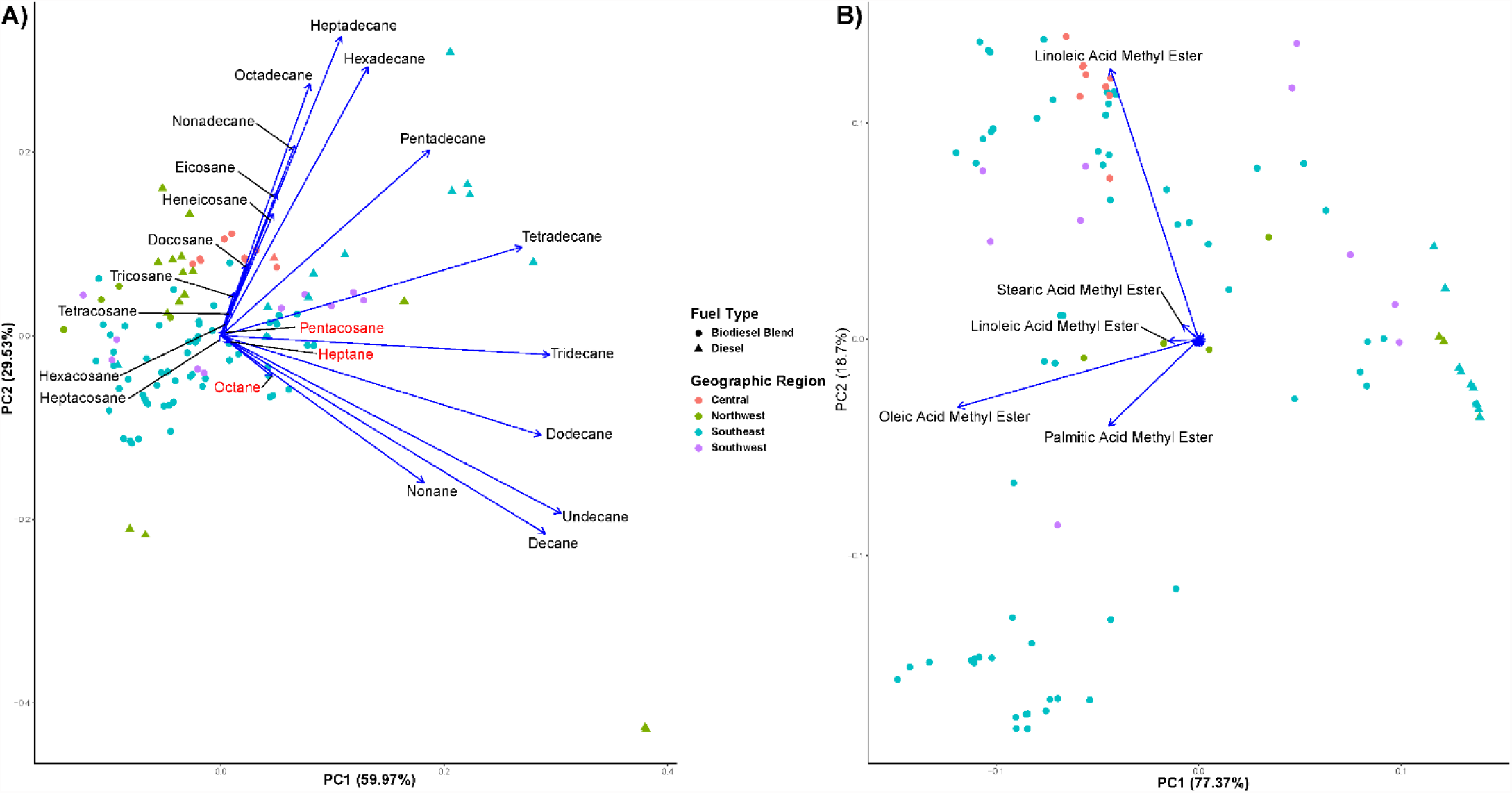
PCA ordination of fuel samples based on n-alkane composition. Biodiesel (circles) and diesel (triangles) fuel samples from different geographical areas in the continental U.S. are represented with different colors with Central (Orange), Northwest (Green), Southeast (Blue), and Southwest (Purple). Blue vectors represent the different n-alkanes and their contribution to the separation of the fuel samples in this ordination space. **B**. Blue vectors represent the different FAME components and their contribution to the separation of the fuel samples in this ordination space. Many FAME components had low eigenvectors in this PCA, and their chemical names are excluded for simplicity; however, their vectors remain in this PCA to give a better overview of how fuel samples are separated.

In contrast to n-alkanes in which fuel region appeared to primarily distinguish fuel composition, fuel type (biodiesel blend or diesel) primarily separated samples, with diesel fuels clustering tightly together when visualized by PCA. The biggest drivers of FAME variability were linoleic, oleic, and palmitic acid methyl esters concentrations and to a lesser extent steric and linoleic acid methyl ester (FIG 1B).

### Microbial Community Analysis of Contaminated Fuels

A total of 150 samples produced 3,834,984 sequence reads for 16S and 523,852 sequence reads for 18S, resulting in an average of 25,567 reads per sample for 16S and 3492 for 18S. Bacterial communities were more diverse than the fungi in contaminated fuels based on the Faith diversity index, where Bacteria had a Faith PD median of 14 in biodiesel samples and 10 in diesel samples, and fungal samples had a Faith PD median of 1.4 in biodiesel and 2.0 in diesel (Supplemental Fig. 1). Some exceptions were detected that appeared to be a bacterial “bloom”, where bacterial communities were composed of more than 80% of the bacterial families *Acetobacteriaceae* and *Moraxellaceae* (FIG 2). Fuels contaminated with *Acetobacteriaceae* were not constrained by geography as these contamination events occurred in both the Northwest and Southeast U.S. Blooms where *Moraxcellaceae* were the most abundant population were only observed in fuel samples from the Southeast U.S.

**FIG 2.**
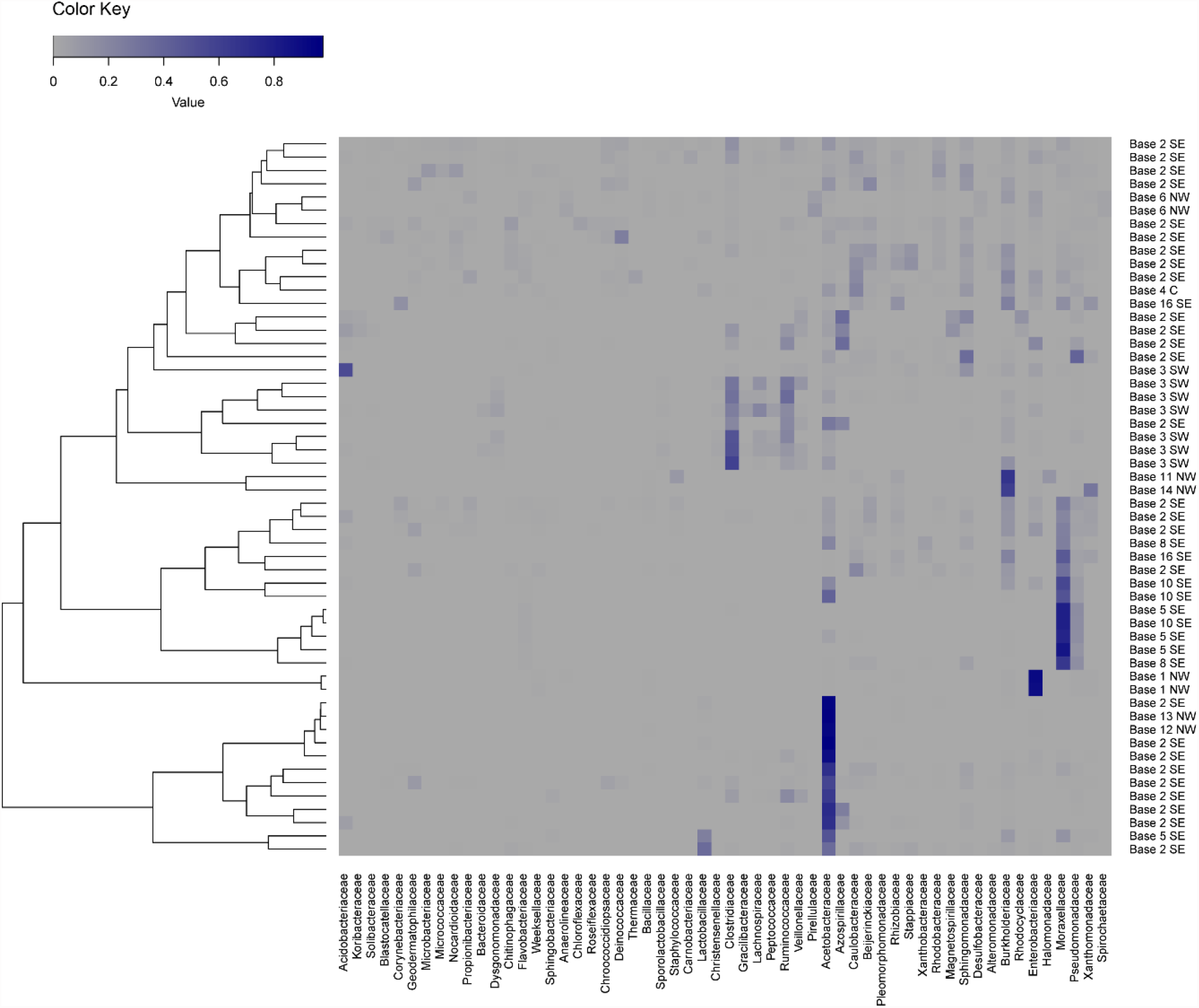
Heatmap representing the relative abundance of bacterial populations in microbial communities from contaminated fuels. Relative abundance of 100% is represented by dark blue, while gray represents 0% of the reads. Samples are sorted by the dendrogram on the left y-axis, which was based on bacterial population similarity. Bacterial families are denoted along the x-axis and sample names are denoted on the right y-axis.

In contrast to the diversity observed among the bacteria, the fungal populations present in microbial communities from contaminated fuels were primarily comprised of a single family, the *Trichocomaceae*. While the *Trichocomaceae* were usually the most abundant family in both diesel and biodiesel blends sequenced, samples within the Southeast U.S. primarily contained members of the family *Debaryomycetaceae* (FIG 3).

**FIG 3.**
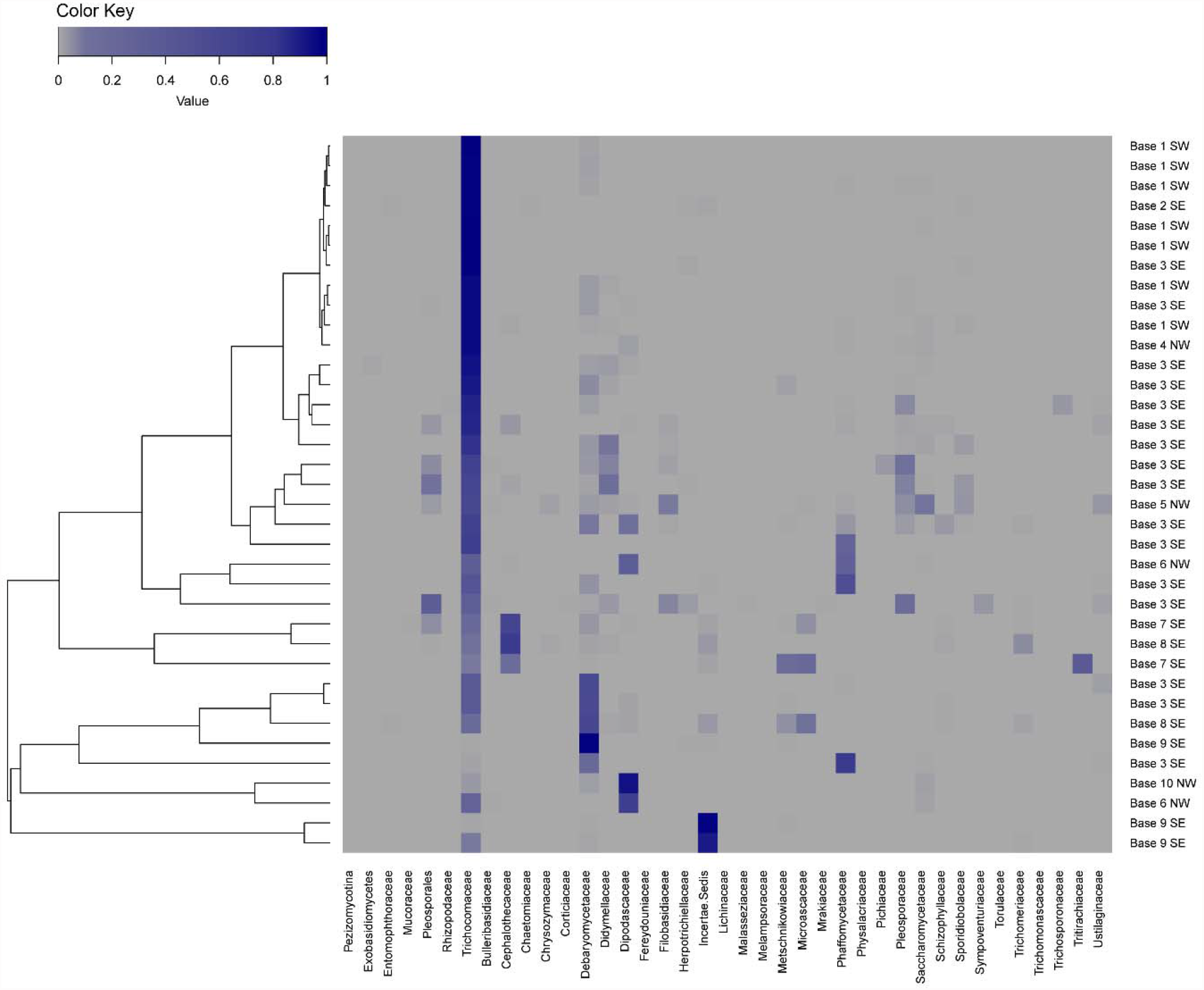
Heatmap representing the relative abundance of fungal populations in microbial communities from contaminated fuels. Relative abundance of 100% is represented by dark blue, while gray represents 0% of the reads. Samples are sorted by the dendrogram on the left y-axis, which was based on fungal population similarity. Fungal families are denoted along the x-axis and sample names are denoted on the right y-axis.

### Redundancy Analysis Correlating Fuel Components to Taxonomy

Redundancy analyses (RDA) were conducted to extract and summarize the variation in the microbial taxonomic data (both bacterial and fungal) and derive correlations with the fuel composition. The bacterial family *Acetobacteriaceae* were positively correlated to fuels containing more long chain n-alkanes including eicosane, tricosane, and tetracosane (FIG 4). The *Acetobacteriaceae* were also positively correlated with increases in linoleic acid methyl ester. In contrast, the bacterial family *Moraxellaceae* was positively correlated with fuels containing shorter n-alkanes and higher concentrations of myrisoleic acid methyl ester. The RDA model from the bacterial taxa and fuel composition had a R^2^ value of 0.33 with p=0.003. Fuel compounds found to significantly impact microbial communities were used in the bacterial RDA using an adonis test on the Hellinger transformed taxonomic data can be seen in Table 1.

**FIG 4.**
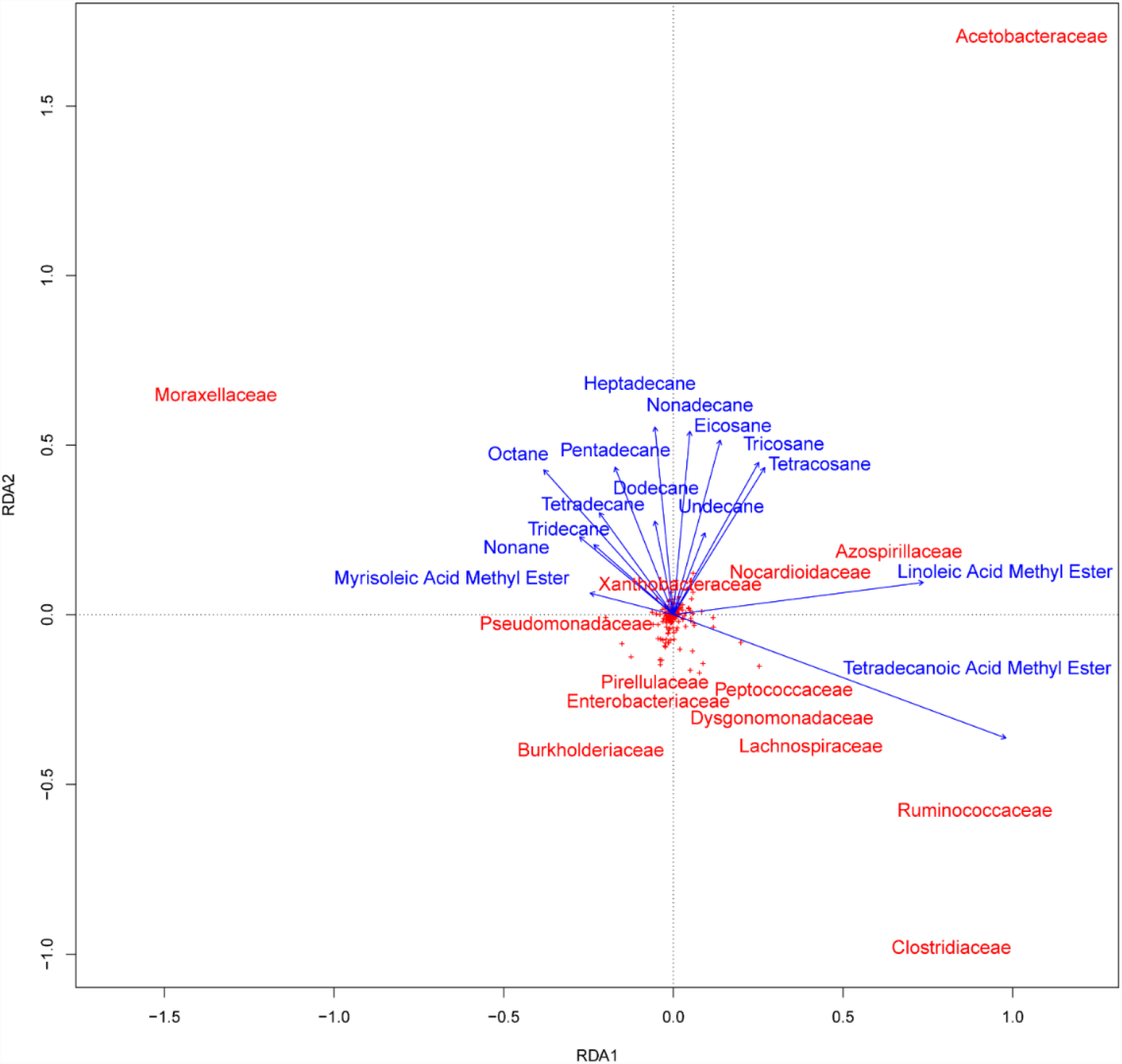
Redundancy analysis (RDA) of the Hellinger-transformed bacterial populations constrained by the fuel composition. Bacterial families are denoted in red text. In this RDA the red crosses are bacterial families that still contribute to this ordination space, but not specifically named due to their central clustering indicating a low correlation coefficient. Fuel component variables are represented by blue vectors. Blue vectors lengths indicate the relative weight of a fuel component in the ordination.

**Table 1.** Forward selection of significant (Pr(>F) ≤ 0.05) fuel variables based on an adonis permutational multivariate analysis of variance on the bacterial Hellinger transformed taxonomic data.

| Fuel Component | Mean Squares | F. Model | R<sup>2</sup> | Pr(>F) |
| --- | --- | --- | --- | --- |
| **Octane** | 0.656 | 4.076 | 0.042 | 0.001 |
| **Nonane** | 0.505 | 3.137 | 0.032 | 0.003 |
| **Decane** | 0.297 | 1.849 | 0.019 | 0.045 |
| **Dodecane** | 0.307 | 1.907 | 0.020 | 0.044 |
| **Tridecane** | 0.342 | 2.121 | 0.021 | 0.041 |
| **Tetradecane** | 0.617 | 3.835 | 0.040 | 0.001 |
| **Pentadecane** | 0.313 | 1.946 | 0.020 | 0.050 |
| **Hexadecane** | 0.389 | 2.415 | 0.025 | 0.019 |
| **Docosane** | 0.466 | 2.896 | 0.030 | 0.005 |
| **Tetracosane** | 0.329 | 2.040 | 0.021 | 0.039 |
| **Pentacosane** | 0.366 | 2.274 | 0.024 | 0.020 |
| **Hexacosane** | 0.489 | 3.033 | 0.031 | 0.004 |
| **Myrisoleic Acid ME** | 0.328 | 2.035 | 0.021 | 0.025 |
| **Tetradecanoic Acid ME** | 0.365 | 2.264 | 0.023 | 0.018 |
| **Linoleic Acid ME** | 0.519 | 3.222 | 0.033 | 0.002 |

An RDA was also generated to identify how fungal communities may be correlated with fuel composition (FIG 5). The abundant fungal family *Trichocomaceae* was positively correlated with fuels containing higher concentrations of FAME. *Debaryomycetaceae* was found to be abundant in fuels containing higher concentrations of pentadecanoic and oleic acid methyl acids. The *Debaryomycetaceae* were also found in fuels containing more mid-chain n-alkanes (*e*.*g*. pentadecane and hexadecane). The fungal RDA had a R^2^ value of 0.45 with p = 0.005. Fuel compounds found to significantly impact fungal communities used in the fungal RDA determined via a permutational multivariate analysis of variance using distance matrices on the Hellinger transformed taxonomic data can be seen in Table 2. The fungal family *Trichocomaceae* had no correlation with the FAME pentadecanoic acid methyl ester which was tested later as a sole carbon and energy source for the representative isolate *Paecilomyces* AF001 for growth.

**FIG 5.**
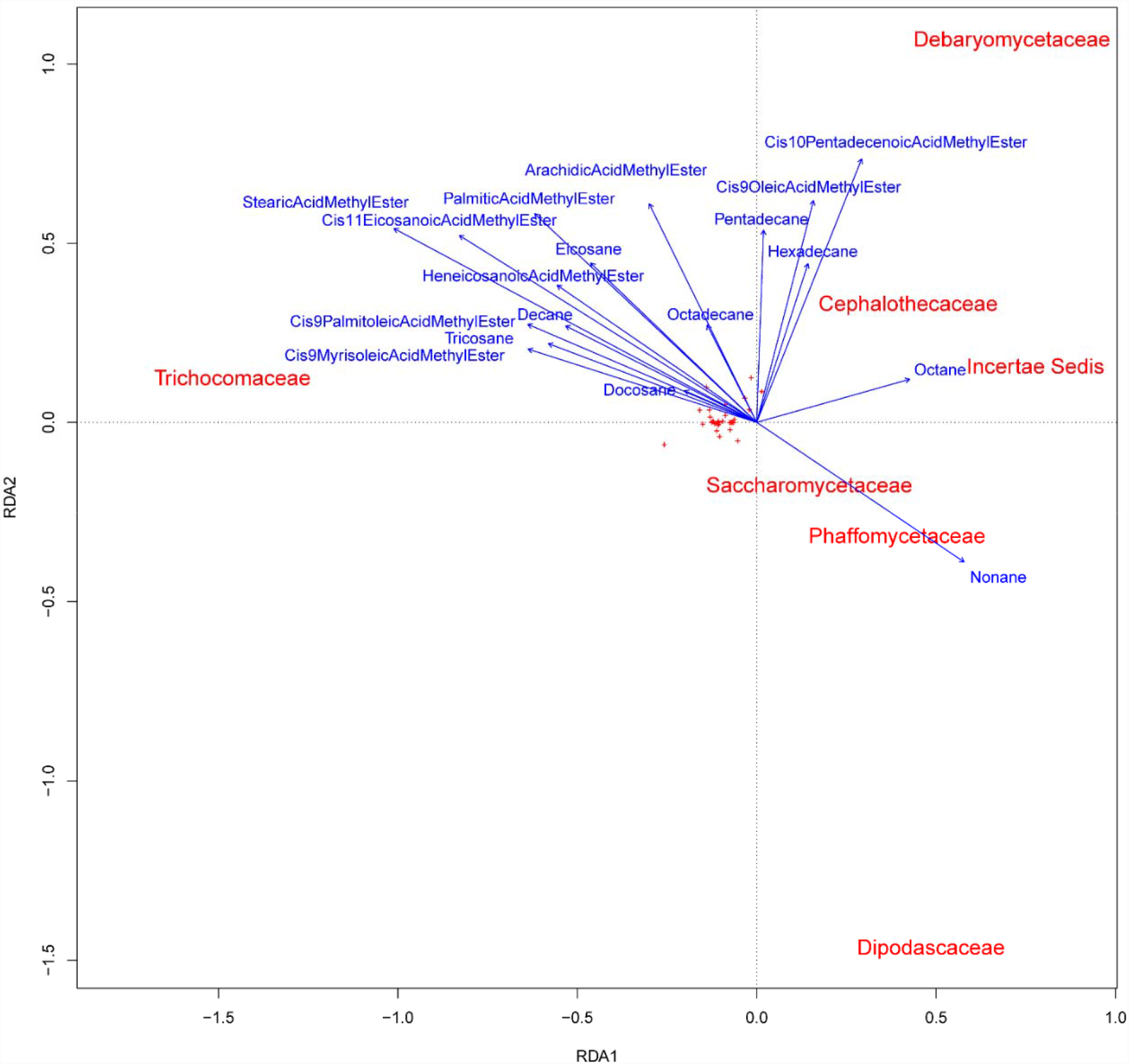
Redundancy analysis (RDA) of the Hellinger-transformed fungal communities constrained by the fuel composition. Fungal families are denoted as red text. In this RDA the red crosses are fungal families that still contribute to this ordination space, but not specifically named due to their central clustering indicating a low correlation coefficient. Fuel component variables are represented by blue vectors. Blue vectors lengths indicate the relative weight of a fuel component in the ordination.

**Table 2.**
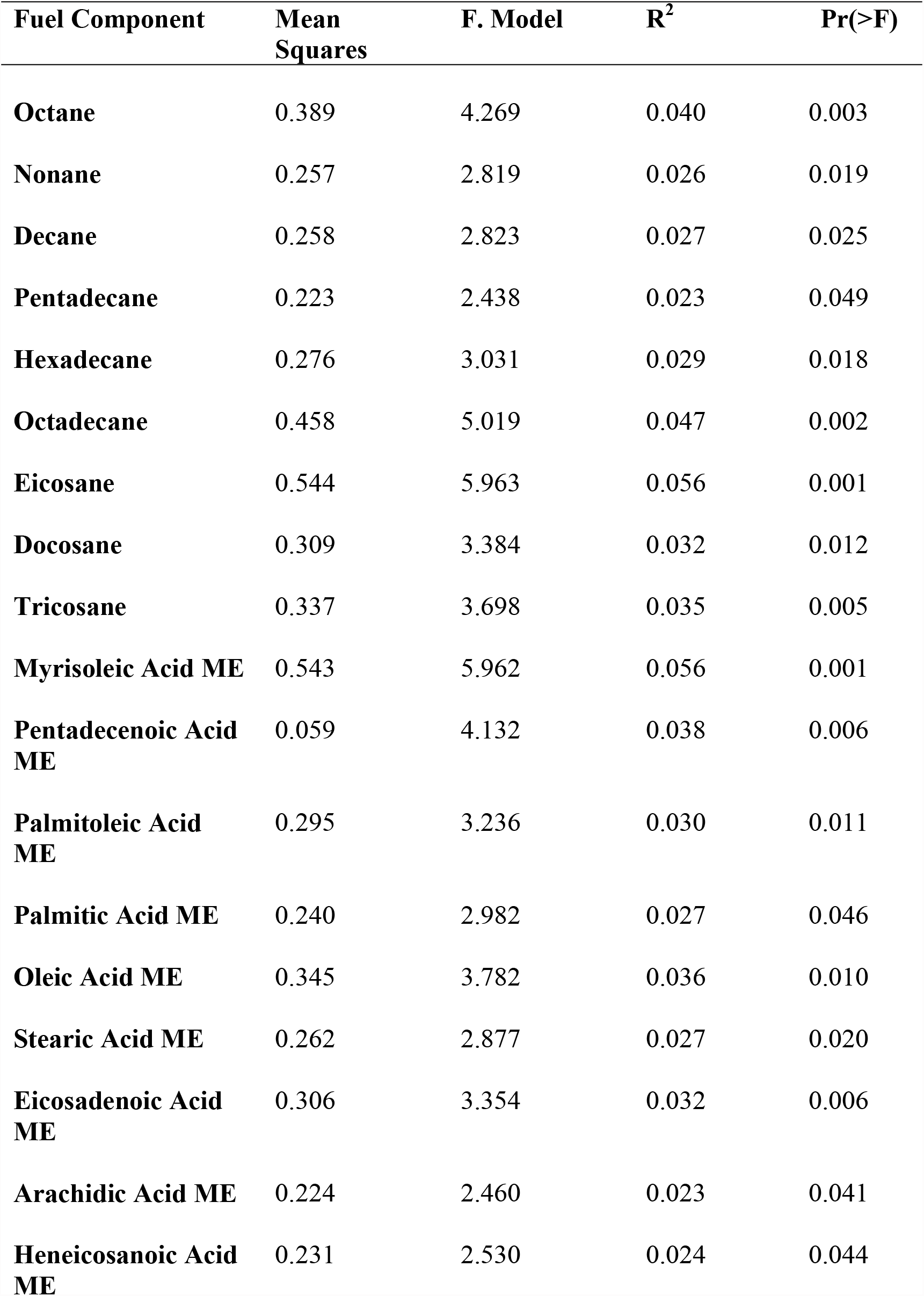
Forward selection of significant (Pr(>F) ≤ 0.05) fuel variables based on an adonis permutational multivariate analysis of variance on the fungal Hellinger transformed taxonomic data

### Physiological Characterization of Isolates to Test Correlations Predicted from Fungal RDA

The RDA identified many correlations between both bacterial and fungal communities and fuel composition. Two isolates representative of abundant fungal genera from contaminated fuels were used to test correlations identified from via RDA (13). The filamentous fungus *Paecilomyces* was able to grow successfully on palmitoleic acid methyl ester as a sole carbon and energy source (strong correlation); however, it was unable to grow using pentadecanoic acid methyl ester (no correlation). *Wickerhamomyces*, which had a strong correlation with both palmitoleic and pentadecanoic acid methyl esters, was able to use them both as sole carbon and energy sources (FIG 6).

**FIG 6.**
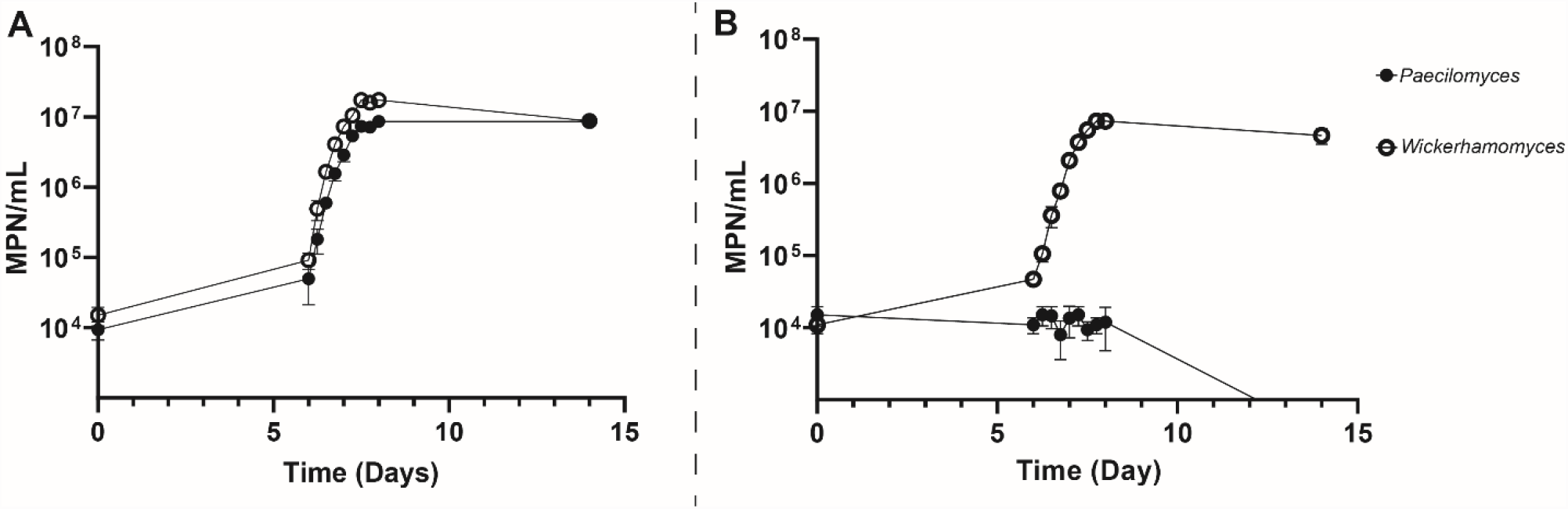
Growth of the fungal isolates *Paecilomyces* AF001 (Filled circles; *Trichocomaceae*) and *Wickerhamomyces* SE3 (Open circles; *Debaryomycetaceae*) palmitoleic acid methyl ester as a sole carbon and energy source. Error bars represent standard deviation for mean *Paecilomyces* AF001 or *Wickerhamomyces* SE3 (n=3). **B:** Growth of *Paecilomyces* AF001 (Filled circles; *Trichocomaceae*) and *Wickerhamomyces* SE3 (Open circles; *Debaryomycetaceae*) on pentadecanoic acid methyl ester as a sole carbon and energy source. Error bars represent standard deviation for mean *Paecilomyces* AF001 or *Wickerhamomyces* SE3 (biological replicates n=3).

## Discussion

The composition of microbial communities can be influenced by numerous factors including the availability of oxidizable substrates that can be differentially consumed (28). We considered this type of selection true in fuel storage tanks where the composition of fuel components may play a significant role in dictating which microorganisms may be prevalent. Correlations were identified between the chemical composition of both B20 biodiesel and diesel fuels and their resident microbial communities. Two positive correlations were between members of the fungal families *Trichocomaceae* and *Debaryomycetaceae* and palmitoleic and pentadecanoic acid methyl esters, respectively. Two representative fungi, *Paecilomyces* AF001 and *Wickerhamomyces* SE3, previously isolated from contaminated fuel storage tanks, were then used to test the correlations. Growth studies with palmitoleic and pentadecanoic acid methyl esters as a sole carbon and energy source verified the link predicted between these abundant members the fuels they inhabited.

Regional differences in fuel composition were identified via GC-MS, in addition to differences more clearly attributable to the type of fuel analyzed (B20 biodiesel vs. ULSD). Most of the differences in n-alkanes of biodiesel and diesel fuels were attributed to the presence of higher chain alkanes (C15-24) vs. lower chain alkanes (C9-C14, Fig 3). Most biodiesel fuel samples were similar to one another, with smaller differences in their n-alkane composition compared to the diesel fuels. As predicted, diesel fuels had more similar FAME profiles, likely because FAME was in much lower abundance compared to the B20 biodiesel blends. Among the B20 biodiesel fuels, the variation in FAME composition was largely based on the concentration of linoleic, palmitic, oleic, stearic, and linolelaidic acid methyl esters. When different feedstocks are used, the resulting biodiesel will have different total percent weights of these FAME (15).

The fuels analyzed in this study were chemically diverse, exhibited differences based on region, but were also correlated with the composition of their microbial contaminants.

The bacterial populations found in contaminated fuels were much more diverse in terms of richness than the fungal populations (Supplementary FIG 1). However, there were some exceptions as some contaminated fuels contained higher relative abundances of bacteria from the families *Acetobacteriaceae* and *Moraxellaceae*. Both bacterial families have been implicated in diesel and biodiesel contamination. Additionally, their metabolism of fuel components produces organic acids, causing increased rates of corrosion (12, 29).

Fungal populations were not as richly diverse as the bacteria in the contaminated fuels. Fungal contamination in sampled fuels was largely comprised of members of the family *Trichocomaceae*. Although the *Trichocomaceae* are ubiquitous in nature and normally associated with the spoilage of fruit juices and foods, they have also been linked to biodiesel degradation and MIC (13, 30). Members of the fungal family *Debaryomycetaceae* were also a primary contaminant in fuels although to a lower extent than *Trichocomaceae*. Members of this yeast family are used in fermentation processes in the food industry but have also been linked to fuel degradation and MIC (12, 13, 31).

Different microbial populations in fuels can have different fouling properties, different metabolic processes, and pose different risks to infrastructure as they metabolize fuels (32, 33). If fuel composition selected for distinct microbial community assemblages, we expected correlations between fuel composition and the bacterial or fungal populations found in contaminated fuel. Members of the bacterial family *Acetobacteraceae* were positively correlated to fuels that contained higher alkane chain lengths than others as well as with linoleic acid methyl ester. Some bacteria, such as *Pseudomonas* and *Acinetobacter*, primarily degrade alkanes with higher chain lengths (C19+) (22, 34). Although it is still not clearly understood which hydrocarbon substrates *Acetobacteriaceae* may preferentially use, they are commonly found in contaminated diesel and biodiesel fuels (12, 35, 36). Their correlation with fuels containing longer chain alkanes (C19-24) would suggest that they are able to use these alkanes, representing a potential niche other bacteria or fungi cannot occupy. Additionally, the bacterial family *Moraxellaceae* was positively correlated with smaller alkane chains (C8-14) and myrisoleic acid methyl ester (Fig 5). Members of this bacterial family have also been observed in contaminated fuels; however, like the *Acetobacteraceae*, little is known about which fuel compounds they specifically metabolize (37, 38).

Correlations were also found between the abundance of fungal populations and certain fuel components. *Trichocomaceae* was the prominent fungal population in contaminated fuels and correlated with fuels containing more fatty acid methyl esters including palmitoleic acid methyl ester. Members of the yeast family *Debaryomycetaceae* were more abundant in fuels containing higher concentrations of Cis-10-Pentadecenoic Acid Methyl Esters and Cis-9-Oleic Acid Methyl Esters (Fig 3.6). The correlations of these two fungal families suggested that they may have differential capabilities in degrading these fuel components. Both members of the *Trichocomaceae* and *Debaryomycetaceae* have been implicated in contamination of diesel and biodiesel fuels previously and appear to be a common fuel contaminant across the continental U.S. when fungal contamination is present (12, 39). Representative organisms from both fungal families have also recently been implicated in corrosion when grown on B20 biodiesel (13).

The predicted difference in fuel component preference between the *Trichocomaceae* and *Debaryomycetaceae* was directly tested *in vitro* using two representative taxa, *Paecilomyces* AF001 and *Wickerhamomyces* SE3, previously isolated from contaminated B20 biodiesel (13). *Wickerhamomyces* SE3 grew well on both palmitoleic and pentadecanoic acid methyl ester, but *Paecilomyces* could only grow on palmitoleic acid methyl ester (Fig 7). Both FAME compounds contain one degree of unsaturation, but palmitoleic acid methyl ester has an even number carbon chain (C16) while pentadecanoic acid methyl ester has an odd number carbon chain (C15). Even though both have one degree of saturation, the double bond in palmitoleic acid methyl ester is at C9, while pentadecanoic acid methyl ester has a double bond at C10, an odd and an even numbered carbon in the fatty acid chains. Not all organisms are able to metabolize odd numbered fatty acids and a more diverse set of enzymes are needed to metabolize the double bonds in the fatty acid chain (40). Odd numbered bonds are typically activated by isomerases while the double bonds on even numbered chains must be activated by isomerases and reductases, which *Paecilomyces* may not have. This would explain why it was not able to grow with pentadecanoic acid methyl ester as a sole carbon and energy source.

The availability of different carbon substrates can influence microbial community composition and differences have been observed in diesel and biodiesel fuels as well (41–45). Correlations between both bacterial and fungal populations in contaminated fuels and the composition of FAME and alkanes in those fuels suggest that fuel composition influences the composition of contaminating microorganisms. A few of these distinct correlations were supported by growth studies with two representative fungi. While carbon substrates in an environment can drive community structure, it is important to note that this is not the only driver. Biotic and abiotic factors such as interactions between microorganisms, the availability of nutrients like phosphorous and nitrogen, or the amount of water present in a contaminated fuel tank can also influence which microorganisms will proliferate. Our findings described here link fuel composition to the microorganisms found in contaminated storage tanks, providing insight as to which feedstocks are best to use when producing biodiesel. For instance, limiting the presence of certain FAMEs, such as those readily consumed by *Paecilomyces* may strongly control biocontamination in tanks. Further, identifying fuel components that encourage the growth of microorganisms that do not cause pervasive biofouling or corrosion would be useful in developing potentially probiotic fuel communities.

## Materials and Methods

### Fuel Tank Sampling Protocol

Fuel was collected from 17 sites across the continental United States. Site selection was contingent upon obtaining access to contaminated fuel storage tanks. A map containing an overview of general site region and the number of fuel samples taken is described in Supplementary FIG 2 and Supplementary Tables 1-3. Fuel from storage tanks was collected using a stainless-steel Bacon Bomb (Koehler™ Instrument Petroleum Bacon Bomb Sampler, Fisher Scientific) that was field disinfected using 100% isopropanol prior to insertion into the tanks. Once field sterilized the Bacon Bomb was inserted into the fuel storage tanks and 500 mL of fuel was collected from the bottom of the tanks and transferred into sterile 1L bottles. Immediately after collection, the fuel was filtered through a polyether sulfone (PES) Steritop Filter Unit™ (Millipore) with a nominal pore size of 0.45 μm affixed to the top of a second sterile 1L bottle. The filter was retained for DNA extraction and the fuel was retained for chemical analysis, both described below.

### DNA Extraction, SSU rRNA Gene Library Preparation, and Sequencing

Following fuel filtration, the PES filter was removed and sectioned into four equal quarters using sterile disposable scalpels. Three sections from each sample were individually placed into Zymo Quick-DNA Fecal/Soil Microbe ZR BashingBead™ Lysis Tubes filled with DNA/RNA Shield™ (Zymo Research Co, Irvine, CA, United States). Homogenization and physical lysis of biomass samples was achieved in the field using a custom tube holder attached to a reciprocating saw, agitated at full speed for 1 min. After homogenization, the samples were shipped back to lab, and stored at -20°C until (≤ 3 days) the DNA was extracted via the Zymo Quick-DNA Fecal/Soil Microbe kit (Zymo Research) according to the manufacturer’s protocol.

Small subunit (SSU) rRNA gene libraries were generated by amplifying the extracted DNA using PCR primers that spanned the V4 and V5 hypervariable regions of the 16S ribosomal RNA gene between position 515 and 926 (*Escherichia coli* numbering), which produced a ∼400 bp fragment for Bacteria and Archaea and ∼600 bp fragments for the Eukarya 18S rRNA gene. These primers were selected to amplify a broad distribution of SSU rRNA genes from all three domains of life (44). The forward primer 515F-Y (5’-**GTA AAA CGA CGG CCA GC**CG TGY CAG CMG CCG CGG TAA-3’) contains the M13 forward primer (in bold) fused to the gene-specific forward primer (non-bold). The reverse primer 926R (5’-CCG YCA ATT YMT TTR AGT TT-3’) was unmodified from Parada et al. (46). 5 PRIME HotMasterMix (Quanta Biosciences, Beverly, MA, United States) was used for all library prep PCR reactions with a final reaction volume of 50 μL. Thermocycler conditions included a 3 min initial denaturation at 95 °C followed by 30 cycles of 95 °C for 45 s, 50 °C for 45 s, 68 °C for 90 s, and a final extension of 68 °C for 5 min. Reactions were then purified with Sera-Mag™ paramagnetic beads (MilliporeSigma, St. Louis, MO) at a final concentration of 0.8x. Following purification, 4 μL of PCR product was used in a barcoding reaction affixing a unique 12 bp barcode with a complementary M13 sequence to those found on the forward primer of the initial reaction (Integrated DNA Technology) to each library in 50 μL reactions. A 6-cycle reaction was carried out with identical cycling conditions to the initial amplification. The resulting barcoded PCR products were purified using Sera-Mag™ beads at a final concentration of 0.8 x and eluted in 50 μL of nuclease free water. The cleaned, barcoded PCR products were quantified using the QuBit™ dsDNA HS assay kit (Thermo Fisher Scientific Inc., Waltham, MA, United States) and pooled in equimolar amounts before concentration using an Amicon® Ultra 0.5 mL centrifugal filter with Ultracel-30K membrane (Millipore Sigma, Billerica, MA, United States) to a final volume of 80 μL. To detect and mitigate the effects of reagent contamination, triplicate extraction blanks (DNA extraction with no sample addition) and negative PCR controls (PCR with no template DNA added) were sequenced as well. The pooled prepared libraries were submitted for sequencing on an Illumina MiSeq (Illumina Inc., San Diego, CA, United States) using V2 PE250 chemistry at the University of Oklahoma Consolidated Core Lab.

### GC-MS Quantification and Characterization of FAME and Alkanes in Fuels

Sampled biodiesel and diesel fuels were analyzed to determine the amount and composition of FAME and n-alkanes present. All fuel samples were filtered prior to shipment to the University of Oklahoma to remove biomass. Fuels were analyzed using a Shimadzu GCMS-QP2010 coupled to a AOC-20i autosampler (Shimadzu, Kyoto, Japan). Fuel was diluted prior to injection using GC grade hexane in a 1:10 ratio. All fuel samples were run in triplicate *(i*.*e*. technical replicate). The autosampler injection syringe was rinsed after each 1μL injection with 100% ethanol three times. After the ethanol rinse the syringe was rinsed once with the diluted fuel sample prior to the injection. A 1:10 split injection was used, representing a final dilution of all fuel samples of 1:100.

A 30.0 m Rtx-5MS column (Restek) with a nominal thickness and diameter of 0.25 μm was used. High purity helium at a linear velocity of 36.8 cm per second was used as the carrier gas. GC injection temperature was set at 300°C while the initial column temperature was 40.0 °C. Column temperature was held at 40.0 °C for 1.5 minutes upon injection. After an initial hold the temperature of the column increased by 10 °C per minute to a final temperature of 300 °C, which was held for 2.75 minutes. Mass spectra were analyzed in scan mode with event times of 0.25 seconds with a scan speed of 2000 and a solvent cut time of 2.75 min, an ion source temperature of 200 °C and an interface temperature of 300 °C. Spectra between m/z 35 and 500 were used for all downstream analyses.

External standards were obtained for FAME (Supelco 37 FAME Mix) and n-alkanes (C7-C40 Saturated Alkanes Standard) to quantify the concentration of fuel components in all samples (in parts per million, or ppm). A three-point standard curve for each reference compound was generated. All standards were diluted in hexane in ratios of 1:50, 1:25, and 1:10. The limit of detection of reference standards was 0.2 ppm. Ion intensities in each fuel sample were then compared to the generated standard curves to produce concentration estimates of FAME and n-alkane components.

### Analysis of SSU rRNA Gene Sequencing Libraries

Small subunit rRNA gene analyses were carried out in QIIME2 version 2019.10.0 (47). Briefly, barcodes from the sequences were extracted using fastx truncate in the usearch64 software package, and samples were demultiplexed prior to operational taxonomic unit (OTU) clustering. Chimeric sequences and singletons were filtered out prior to clustering using the USEARCH64 reference database. Representative sequences for each OTU were assigned a taxonomic identity with the QIIME2 sklearn classifier against the SILVA 132 database clustered to a 97% identity (48, 49). Following this, sequences were separated to generate feature tables that contained separated 16S and 18S rRNA gene data. After OTU generation and sequence alignment using MAFFT, 16S rRNA and 18S rRNA phylogenetic trees were generated using FastTree 2 (50). Datasets were rarefied to account for variable library size between samples to 1000 sequences per sample for bacteria based on rarefaction curves and 250 sequences per sample for Eukarya due to the lower sequence and diversity in these samples (54). These data were then converted to BIOM files and exported for further analyses in R (51).

### Redundancy Analyses (RDA)

Fuel and taxonomic relative abundance data were used to generate redundancy analyses to determine how fuel components correlated to microbial community composition. Taxonomic data generated in QIIME 2 were imported into R and transformed using the Hellinger method to account for the sparsity of amplicon data (55). Transformed taxonomy data were then used in a forward selection of fuel variables, carried out using a permutational multivariate analysis of variance using distance matrices test in R using the function ‘adonis’ within the vegan package to determine which fuel components were significantly correlated (p<0.05) with community composition. An RDA was then generated by using the transformed community data and those fuel components that were significantly correlated to microbial community composition.

### Validation of Fungal RDA Model

FAME components that were strongly correlated with *Trichocomaceae* and *Debaryomycetaceae* from the RDA model were used to determine how well representative organisms isolated from contaminated fuels could grow on individual FAMEs as a sole carbon and energy substrate. Prior work from our lab yielded the fungal isolates *Paecilomyces* AF001 and *Wickerhamomyces* SE3 which are representative of the fungal families *Trichocomaceae* and *Debaryomycetaceae* respectively (13). A spore suspension of *Paecilomyces* AF001 and a cell suspension of *Wickerhamomyces* SE3 were used to inoculate a 1:20 FAME substrate in artificial sump water (ASW, per L: 0.015g NaCl, 0.035g NaF, 0.02g CaCl_2_, 0.018g KNO_3_, 0.01g Na_2_SO_4_, 0.015g (NH_4_)_2_SO_4_, and 0.017g K_2_HPO_4_) (51).

To prepare a spore suspension, a glycerol stock of *Paecilomyces* AF001 was struck onto Hestrin Schramm (HS) agar medium (per L: 20g glucose, 5g yeast extract, 5g peptone, 2.7g Na_2_HPO_4_, 1.15g citric acid, 7.5g Agar; pH adjusted to 6.0 with diluted HCl or NaOH) and incubated at 25 °C for 7 days (51) After 7 days 5mL of phosphate buffered saline was added overtop the HS agar containing hyphal growth of *Paecilomyces* AF001 and an inoculating loop was used to scrape off the fungal growth to produce a suspension of cells and spores. The fungal suspension was then collected from the agar plate and filtered through a 10 μm nominal pore size polyether sulfone filter to separate spores from the hyphal biomass. The filtrate containing spores was then centrifuged at 10,000 x RCF for 1 minute. The supernatant was decanted, and sterile PBS was added back to the spore pellet and vortexed to resuspend the spores. This wash step was repeated for a total of three washes. *Paecilomyces* spore concentrations were determined using a Petroff-Hausser counting chamber and then diluted in ASW to adjust the final inoculum concentration to 1×10^4^ spores/mL.

To produce a suspension of yeast cells *Wickerhamomyces* SE3 was grown in HS broth for 48 hours and centrifuged at 10,000 RCF to pellet cell mass. Following centrifugation, the supernatant was decanted, sterile PBS was added to the cell pellet, and finally vortexed to resuspend the cells. This was step was repeated for a total of three washes. Cell concentrations were again determined using a Petroff-Hausser counting chamber and diluted to adjust the initial inoculum concentration to 1×10^4^ cells/mL.

*Paecilomyces* AF001 and *Wickerhamomyces* SE3 were grown on palmitoleic and pentadecanoic acid methyl esters (Fisher Scientific, Waltham, MA) to determine their capacity on using these substrates for growth. A total volume of 5 mL 1:20 FAME and artificial sump water mixture was made and inoculated with 10^4^ spores or cells per mL for T=0. Growth was measured by destructively sampling triplicate test tubes at time points 0, 6, 6.25, 6.5, 6.75, 7, 7.25, 7.5, 7.75, 8, and 14 days at room temperature and generating MPNs.

### Statistical Analyses and Data Visualization

Statistical analyses and figure generation was carried out in R version 3.3.3 and GraphPad Prism 8.3.0. Significant differences from fuel components that contributed to microbial community structure were determined via a permutational multivariate analysis of variance using distance matrices on the Hellinger transformed taxonomic data.

## Acknowledgements

We acknowledge the men and women of the US Air Force and Civilian Personnel at US Air Force bases; their cooperation and assistance were critical to this research. Additionally, we would like to thank Dr. Emily Junkins for insightful comments during the development of this manuscript. This work was supported by the Air Force Research Laboratory Biological Materials and Processing Research Team, Materials and Manufacturing Directorate and the U.S. Department of Defense Office of Corrosion Policy & Oversight Technical Corrosion Collaboration grant to B.S.S. (Grant # FA7000-15-2-0001).

## Data Availability Statement

The raw sequence data generated for this study can be found in the Sequence Read Archive under the accession numbers SRR5826605–SRR5826609.

